# Two mammalian-like cysteine dioxygenases and a sulfite exporter play important roles in sulfur homeostasis and virulence of a human pathogen

**DOI:** 10.64898/2026.04.22.720190

**Authors:** Argha Saha, Denisa Petráčková, Jana Holubová, Branislav Večerek

## Abstract

*Bordetella pertussis* is a strictly human, re-emerging respiratory pathogen and the causative agent of whooping cough. Through its adaptation to humans, *B. pertussis* decayed or lost genes of the sulfate assimilation pathway and, consequently, must obtain cysteine from the host. Previously, we showed that the sulfur metabolism of *B. pertussis* is substantially rewired during infection of human macrophages. Here, we investigated the role of several cysteine metabolism- and transport-related genes in the fitness and virulence of this pathogen. We show that excess cysteine strongly induces the expression of genes encoding two cysteine dioxygenases, BP2871 and BP3011, and a putative sulfite exporter, BP2808. The mutant lacking both cysteine dioxygenase genes exhibits impaired growth *in vitro*, severely reduced secretion of pertussis toxin, and attenuated virulence *in vivo*. We also demonstrate the essential role of the sulfite exporter BP2808 and γ-glutamyl-cysteine synthase BP0598 in the adaptation of *B. pertussis* to stress induced by excess cysteine. Intriguingly, both cysteine dioxygenases contain cysteine near the active site, a feature typical of mammalian enzymes and associated with the capacity to increase CDO activity and stability in response to excess cysteine. We hypothesize that the presence of cysteine represents an evolutionary adaptation that improves the survival of *B. pertussis* within a mammalian host. Overall, our data suggest that sulfur metabolism has been effectively streamlined in *B. pertussis* and plays an important role at the host-pathogen interface.

**Importance:** Sulfur is one of the essential nutrients required by cells for growth and cysteine is central to sulfur metabolism. While most bacteria prefer environmentally available sulfate as their cysteine source, several bacterial pathogens rely on cysteine provided by the host. Here we show that *Bordetella pertussis*, the causative agent of whooping cough, has simplified its sulfur metabolism. Our data suggest that two cysteine dioxygenases and sulfite exporter play key roles in sulfur homeostasis and redox balance. Both dioxygenases enable the pathogen to use cysteine as a source of sulfur and the sulfite exporter removes the toxic byproduct of cysteine conversion. Importantly, lack of cysteine dioxygenase activity leads to aberrant secretion of pertussis toxin, one of the essential virulence factors, resulting in attenuated virulence of the pathogen. We suggest that cysteine auxotrophy can be considered part of an infection strategy that assists *B. pertussis* in adapting to its human host.

## Introduction

Sulfur is one of the essential nutrients required by cells for growth. In most cells, sulfur is incorporated into sulfur-containing amino acids cysteine and methionine, cofactors such as thiamine and biotin, and redox-active compounds such as iron-sulfur clusters and cysteine-based thiols (1). Due to its ability to adopt a wide range of oxidation states, sulfur plays a critical role in maintaining cellular redox balance and resistance to oxidative stress (2, 3). Bacteria sense the concentration of intracellular cysteine, which indicates whether the cells have sufficient levels of sulfur, and respond to sulfur or cysteine limitation by inducing the synthesis of specific transporters that import sulfur-containing compounds. Most bacteria prefer inorganic sulfate as their sulfur source and use the sulfate assimilation pathway to synthesize cysteine (4, 5). Thus, cysteine is central to sulfur metabolism and performs diverse functions that cannot be easily fulfilled by other amino acids, such as forming disulfide bridges that stabilize periplasmic and secreted proteins. Cysteine also provides redox activity, however, excess cysteine can be toxic because it can reduce ferric iron to its ferrous form, which can then interact through the Fenton reaction with hydrogen peroxide, leading to a generation of hydroxyl radicals (6). Therefore, cysteine levels within the bacterial cell must be well balanced. Cysteine dioxygenases (CDOs) are important enzymes that can prevent the harmful accumulation of free cysteine by its irreversible oxidation to cysteine sulfinic acid (7). Importantly, cysteine sulfinic acid is a non-thiol compound that can be further broken down to pyruvate and sulfite. Thus, CDOs not only scavenge redox-active sulfur but also link sulfur transactions with central carbon metabolism. Importantly, cysteine, along with glutamate and glycine, is a precursor of reduced glutathione (GSH), a potent thiol antioxidant that plays an important role in redox balance and resistance to oxidative stress (3). Bacteria can import or synthesize GSH to maintain intracellular redox balance (8). GSH biosynthesis proceeds through two enzymatic reactions (9). In the first and rate-limiting step, cysteine is ligated to glutamate by γ-glutamyl-cysteine synthase. In the second step, glutathione synthetase catalyzes the condensation of the previously formed γ-glutamyl-cysteinyl adduct with glycine, yielding GSH.

*Bordetella pertussis* is a Gram-negative strictly human pathogen of the respiratory tract and the etiological agent of whooping cough (pertussis) (10). This highly contagious disease is particularly severe in infants and remains a major cause of infant mortality and morbidity worldwide, especially in developing countries (11). In the last decades, pertussis incidence has been on the rise in industrialized countries with highly vaccinated populations (12, 13). Although the incidence of pertussis has declined during the COVID-19 pandemic, since 2023 many countries have experienced epidemics of pertussis (14). The global resurgence of pertussis suggests that we need to expand our understanding of physiological processes underlying the pathogenesis of *B. pertussis*, including sulfur acquisition at the host-pathogen interface. As a result of speciation to its human host, several genes of *B. pertussis* involved in the sulfate assimilation pathway have been decayed or lost (Fig. S1). Consequently, *B. pertussis* is auxotrophic for cysteine (15) and must acquire cysteine from the host. Our recent parallel transcriptomic profiling of human THP-1 macrophages infected with *B. pertussis* revealed that upon internalization, *B. pertussis* rewires its sulfur metabolism (16, 17). Specifically, genes involved in the catabolism of cysteine and export of its metabolic products were strongly induced. Among them we identified genes encoding two CDO enzymes and putative cysteine, sulfite and sulfate exporters. Remarkably, the mutant lacking two *cdo* genes, *bp2871* and *bp3011*, displayed significantly reduced cytotoxicity toward THP-1 macrophages, indicating the importance of cysteine homeostasis for pathogenicity of *B. pertussis*.

In this study, we investigate the role of several sulfur metabolism-related genes in the fitness of *B. pertussis*. We report that deletion of two mammalian-like *cdo* genes, *bp2871* and *bp3011*, results in attenuated virulence in the mouse model of infection, most likely due to impaired secretion of the pertussis toxin. We also show that the gene encoding the putative sulfite exporter *bp2808* is essential for sulfur homeostasis in *vitro*. Collectively, our data suggest that sulfur metabolism plays an important role in the physiological fitness and virulence of the re-emerging human pathogen *B. pertussis*.

## Results

### The mutant lacking two *cdo* genes is sensitive to increased levels of cysteine in the absence of glutathione

Excess cysteine can be toxic (6) and thus, cysteine auxotrophy in *B. pertussis* may also be seen as an adaptation mechanism to maintain strict control over cysteine levels. In this context, the absence of cysteine dioxygenases (CDOs), which play an important role in scavenging cysteine, could be potentially harmful to *B. pertussis* cells. Our previous research showed that cytotoxicity toward human THP-1 macrophages exerted by the *Δbp2871Δbp3011* mutant, which lacks the two *cdo* genes, was strongly reduced (16). To test the importance of CDO enzymes for the fitness of *B. pertussis in vitro*, we cultivated the wild-type (wt) strain and the *Δbp2871Δbp3011* mutant in standard SS medium and in SS medium supplemented with cysteine to final concentrations of 1.32 mM or 2.64 mM, corresponding to a 4-fold and 8-fold increase in cysteine concentration, respectively, (Fig. 1A, B and C). The mutant strain did not display a growth defect compared to the wt strain, and with increasing concentrations of cysteine, both strains reached higher optical density (OD) values. This experiment revealed that even in the absence of *cdo* genes, *B. pertussis* can handle increased amounts of cysteine. However, this experiment was performed in standard SS medium, which also contains 0.32 mM GSH, a well-known physiological antioxidant critical for maintaining redox homeostasis (8). To test whether GSH alleviated the toxicity of cysteine, we repeated the experiment in SS medium lacking GSH. Interestingly, in the absence of GSH, the mutant strain exhibited significantly impaired growth in the presence of 1.32 mM cysteine compared to the wt strain (Fig. 1D). This growth defect became more prominent when the mutant strain was grown in the presence of 2.64 mM cysteine (Fig. 1E).Evidently, in the absence of the cysteine-scavenging activity of CDOs, the redox-balancing activity of GSH is critical. GSH is a tripeptide composed of glutamate, cysteine, and glycine, and the first two amino acids are present in SS medium while glycine can be synthesized from glutamate. Nonetheless, we hypothesized that the addition of glycine to the SS medium could favor GSH biosynthesis and thereby rescue the impaired growth of the *cdo* mutant. Therefore, we grew the wt strain and the mutant in SS medium supplemented with 2.64 mM cysteine and 0.3 mM glycine. In line with our hypothesis, the addition of 0.3 mM glycine alleviated the growth defect of the mutant strain (Fig. 1F) demonstrating that GSH can compensate for the absence of both *cdo* genes. Furthermore, this result suggests that *B. pertussis* is capable of *de novo* GSH biosynthesis.

**FIG 1.**
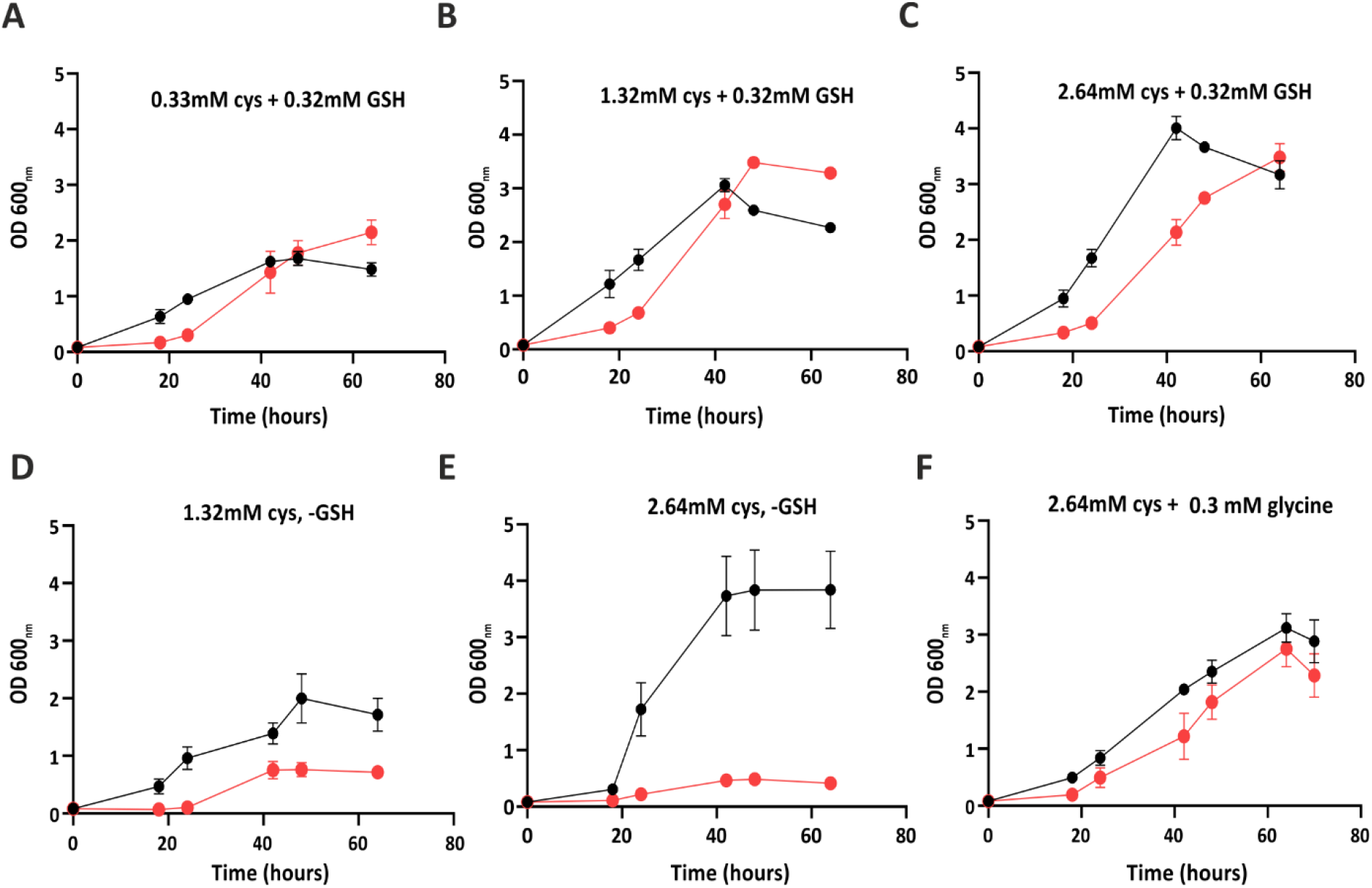
Effect of cysteine, glutathione, and glycine on growth properties of the double *cdo* mutant. The wt strain (black circles) and the Δ*bp3011/bp2871* double *cdo* mutant (red circles) were grown in the standard SS medium in the presence (**A**-**C**) or absence (**D**-**F**) of 0.32 mM GSH. The cysteine concentration in the medium was 0.33 mM (**A**), 1.32 mM (**B** and **D**) and 2.64 mM (**C, E**, and **F**). Cultures shown in panel **F** were additionally supplemented with 0.3 mM glycine. Data represent the mean optical density (OD_600_) ± standard deviation (SD) from three independent biological replicates.

Indeed, our gene homology search identified *bp0598* as a plausible candidate for the γ-glutamyl-cysteine synthase gene *gshA* (Fig. S2), which is required for the first and rate-limiting step in GSH biosynthesis. To verify this assumption, we created a Δ*bp0598* mutant and cultivated it along with the wt strain in SS medium supplemented with 2.64 mM cysteine, either without or with 0.32 mM GSH. In the absence of GSH, the Δ*bp0598* mutant displayed strong growth impairment (Fig. 2A), while in the presence of GSH, the mutant grew similarly to the wt strain (Fig. 2B). These results suggest that *bp0598* is required for the growth of *B. pertussis* in the absence of GSH and most likely represents a *gshA* gene involved in GSH biosynthesis.

**FIG 2.**
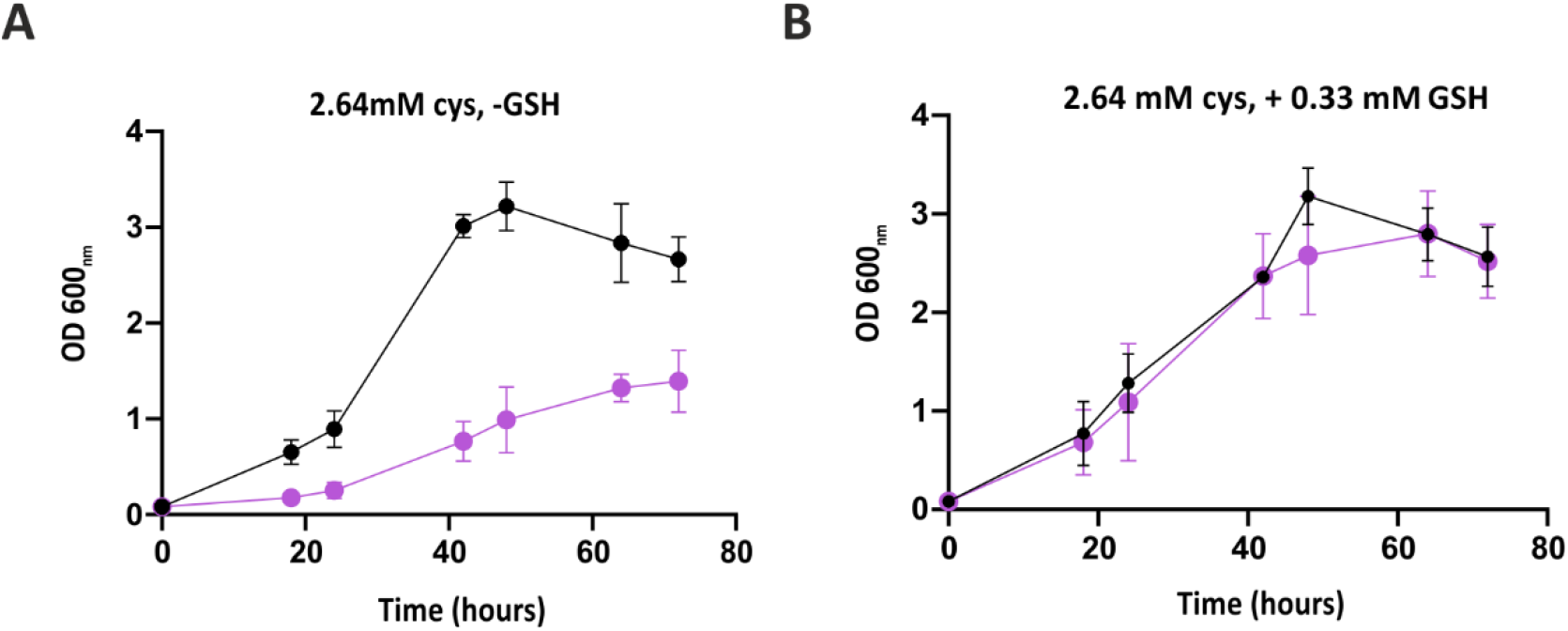
The putative *gshA* gene *bp0598* is required for the growth of *B. pertussis* in the absence of GSH. The wt strain (black circles) and Δ*bp0598* mutant (violet circles) were grown in SS medium supplemented with 2.64 mM cysteine in the absence (**A**) or presence (**B**) of 0.32 mM GSH. Data represent the mean optical density (OD_600_) ± SD from three independent biological replicates.

### Growth phase-dependent transcriptomic analysis indicates changes in the expression of cysteine-related genes

Our dual RNA-seq analysis revealed that, in addition to *cdo* genes *bp2871* and *bp3011*, expression of several cysteine transport-related genes was modulated in intracellular *B. pertussis* cells (16). Therefore, we investigated the expression of both *cdo* genes and genes encoding an ABC transporter (BP1363, a homolog of the cystine importer permease TcyL), a putative cysteine exporter (BP2026, a transporter of the Eam family), a putative sulfoacetate/sulfite exporter (BP2027, a homolog of the TauE exporter), and a putative sulfate exporter (BP2808, an exporter of the YeiH family). The wt cells were cultivated to early (OD_600_ ≈ 0.7-0.8), mid (OD_600_ ≈ 1.1-1.3), and late (OD_600_ ≈ 1.8-2.0) exponential phases of growth, and gene expression was compared between cells grown in SS medium with 2.64 mM cysteine and standard SS medium (control) using quantitative PCR. In the early exponential phase, a significant induction of *bp3011* was observed, while other genes did not show a significant change in expression (Fig. 3A). In the mid exponential phase, *bp3011* was upregulated more than 10-fold, and *bp1363* also showed more than 7-fold induction (Fig. 3B). Finally, in the late exponential phase, *bp1363* was strongly induced with more than a 20-fold increase compared to the control. The expression of the *bp2026* and especially the *bp2027* exporter gene was also significantly induced (Fig. 3C). These experiments indicate that, in contrast to *bp2871*, the expression of the *bp3011* gene is induced when cysteine is present in excess (*i*.*e*., early and mid exponential phases). As anticipated, the expression of the cysteine importer gene *bp1363* is highest in the late exponential phase when cysteine levels are likely reduced, and is accompanied by highly increased expression of the putative sulfite exporter gene *bp2027*.

**FIG 3.**
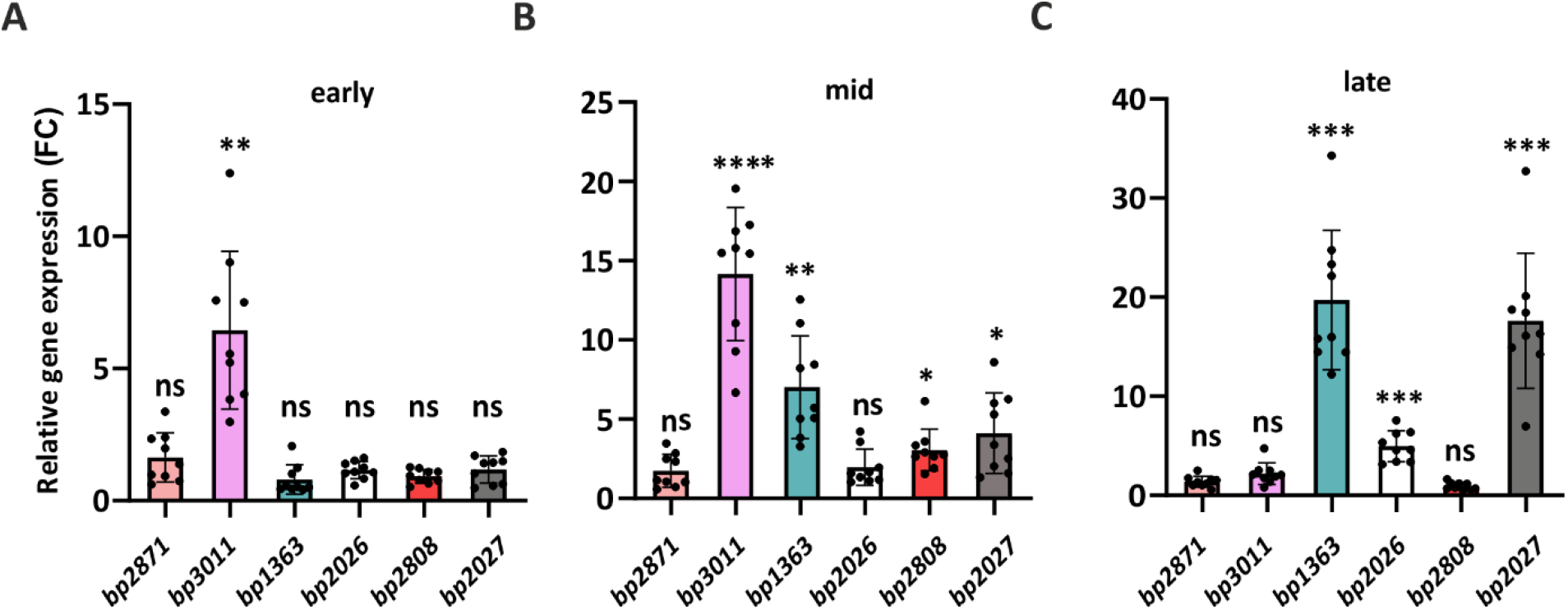
Relative expression of *B. pertussis* genes related to cysteine metabolism. Expression of *bp2871, bp3011, bp1363, bp2026, bp0598*, and *bp2808* genes was determined by quantitative PCR. The wt cells were grown in biological triplicates in standard SS medium (controls) and in SS medium supplemented with 2.64 mM cysteine to early (**A**), mid (**B**) and late (**C**) exponential phases of growth. Gene expression in the presence of 2.64 mM cysteine was compared with that in standard SS medium (set to 1). RT-qPCR analysis of each sample was performed in technical triplicate. The graphs show all individual data points, mean values and error bars representing SDs. Statistical significance was calculated using one-way ANOVA (Dunnett’s multiple comparison test): ns, not significant; (*) *P*-value < 0.05; (**) *P*-value < 0.005; (***) *P*-value < 0.0005; (****) *P*-value < 0.0001.

### The expression of both *cdo* and *yeiH* exporter genes is strongly induced shortly after the addition of excess cysteine

Our data indicated that excess cysteine has only a marginal effect on the expression of the *cdo* gene *bp2871*. This was puzzling, as the expression of the *bp2871* gene was among the most highly induced genes in phagocytosed *B. pertussis* cells (16). Therefore, we asked whether *bp2871* expression is induced only transiently, for example shortly after the addition of cysteine. To investigate the immediate response of *bp2871* and other cysteine export-related genes to excess cysteine, the wt strain was grown in SS medium supplemented with 0.32 mM GSH (as the sole sulfur source) to a mid exponential phase (OD_600_ of approximately 1.0). The culture was then split into three aliquots, which were further incubated in the absence (control) or presence of 0.33 mM and 2.64 mM cysteine for 30 minutes. Total RNA isolated from all cultures was used for quantitative PCR analysis of cysteine metabolism-related genes. Upon addition of 0.33 mM cysteine (the concentration present in standard SS medium), only the expression of *bp2871, bp3011*, and *bp2808* genes was significantly increased by 8-, 10-, and 65-fold, respectively (Fig. 4A). Notably, when the cells were exposed to 2.64 mM cysteine, the *cdo* genes *bp2871* and *bp3011*, as well as the exporter gene *bp2808*, showed a dramatic increase in expression by 50-, 80-, and 140-fold, respectively (Fig. 4B). In contrast, the cysteine importer gene *bp1363* was significantly repressed by elevated levels of cysteine, illustrating the futility of cysteine import in this scenario.

**FIG 4.**
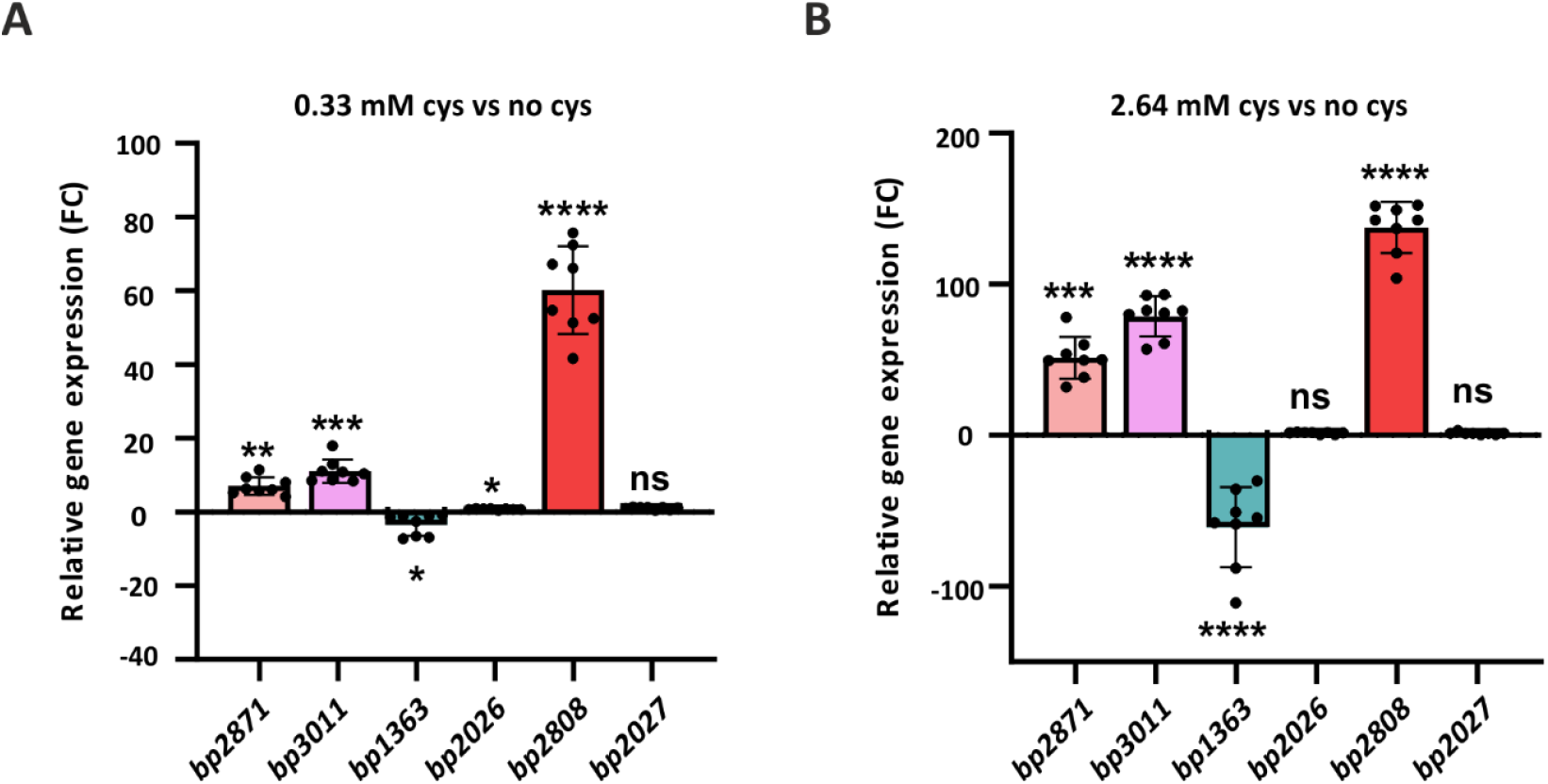
The genes encoding CDOs and the putative sulfate exporter are highly overexpressed after cysteine addition. Expression of *bp2871, bp3011, bp1363, bp2026, bp2027*, and *bp2808* genes was measured in biological triplicates of the wt strain cells harvested 30 min after addition of 0.33 mM (**A**) and 2.64 mM (**B**) cysteine. Gene expression in cells harvested before addition of cysteine was set to 1. RT-qPCR analysis of each sample was performed in technical triplicate. The graphs show all individual data points, mean values, and error bars representing SDs. Statistical significance was calculated using one-way ANOVA (Dunnett’s multiple comparison test): ns, not significant; (**) *P*-value < 0.005; (***) *P*-value < 0.0005; (****) *P*-value < 0.0001.

### The *yeiH* exporter gene *bp2808* plays a significant role in sulfur homeostasis

Our experiments revealed that, in addition to the *cdo* genes *bp2871* and *bp3011*, expression of the exporter gene *bp2808* is also highly induced after the addition of cysteine. BP2808 is annotated as an inner membrane YeiH-like sulfate exporter and shares approximately 40-60% homology with YeiH proteins of other Gram-negative bacteria (Fig. S3). However, an exporter of the YeiH family was recently shown to play an important role in sulfite stress resistance in *S. enterica* (18). To investigate the specificity of the BP2808 exporter, we generated a Δ*bp2808* mutant and cultivated it alongside the wt strain in SS medium and in SS medium supplemented with 0.1 mM, 0.5 mM, and 2 mM sulfate or sulfite. While growth of the Δ*bp2808* mutant in the medium supplemented with sulfate was only temporarily affected compared to the wt strain (Fig. 5A), the impact of sulfite on growth of the mutant was severe. The mutant grew very poorly with 0.1 mM and 0.5 mM sulfite, and in the presence of 2 mM sulfite, it did not grow at all, indicating that the BP2808 transporter is essential for *B. pertussis* cells exposed to excess sulfite (Fig. 5B). Notably, growth of the wt strain was also affected, but to a much lesser extent, as the OD_600_ values in the presence of 2 mM sulfite were slightly reduced compared to those with 0.1 mM sulfite.

**FIG 5.**
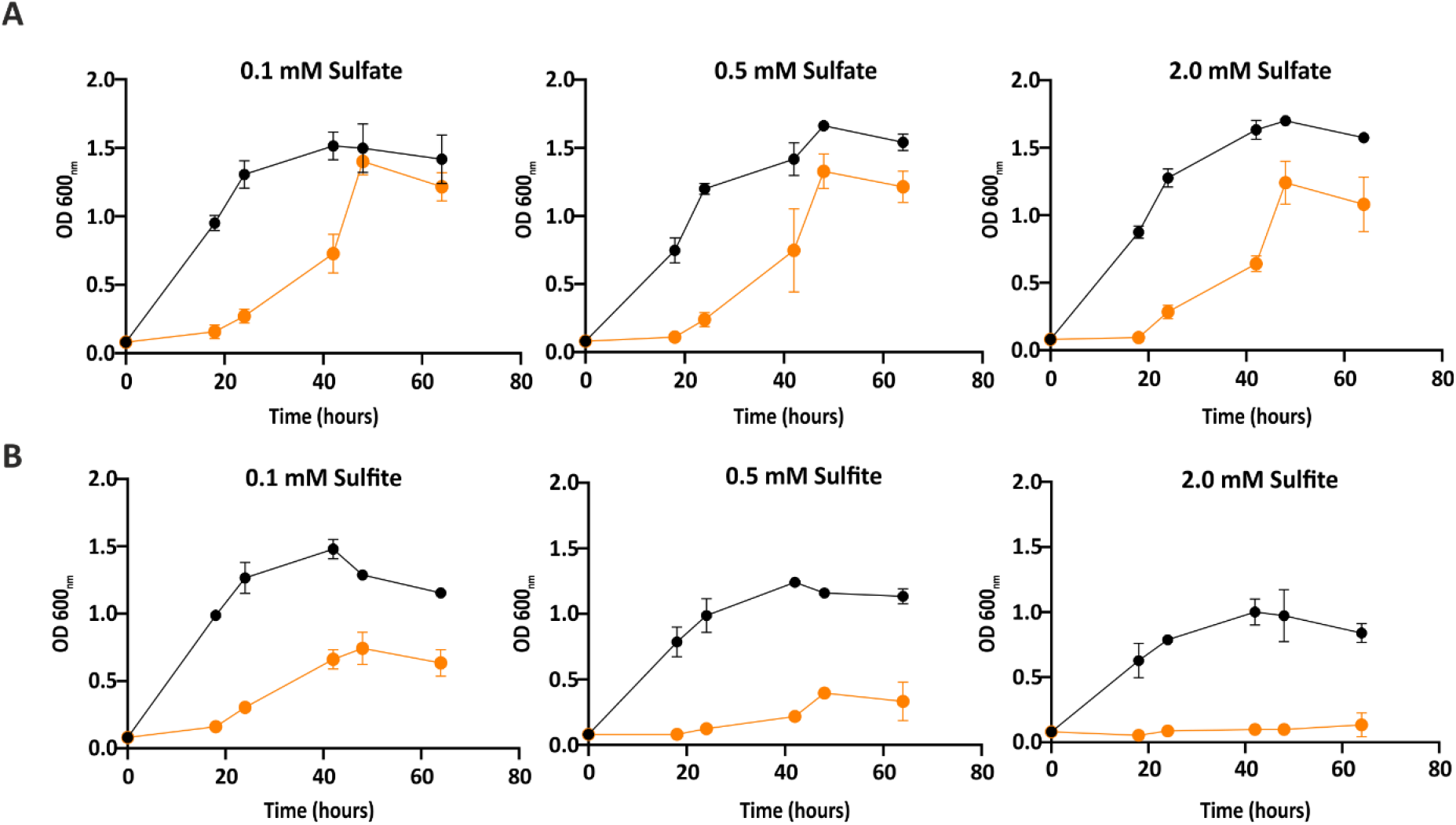
Growth of the Δ*bp2808* mutant is impaired in the presence of sulfite. The wt strain (black circles) and the Δ*bp2808* mutant (orange circles) were grown in standard SS medium supplemented with 0.1 mM, 0.5 mM, and 2 mM sulfate (**A**) or sulfite (**B**). Data represent the mean optical density (OD_600_) ± SD from three independent biological replicates.

### *B. pertussis* appears to favor exporting cysteine metabolites over recycling

Since the production of sulfite and its conversion to sulfate are integral parts of cysteine catabolism, we tested the growth of the Δ*bp2808* mutant in the presence of excess cysteine. The wt strain and the mutant were grown in standard SS medium and in SS medium containing 2.64 mM cysteine. The experiment revealed that growth of the Δ*bp2808* strain is not affected by excess cysteine, although the mutant resumed growth only after a long lag phase compared to the wt strain (Fig. 6A). However, in contrast to the wt strain, the mutant did not benefit from increased concentration of cysteine, as in the presence of 2.64 mM cysteine, the wt strain reached higher OD_600_ values than the mutant (Fig. 6B).

**FIG 6.**
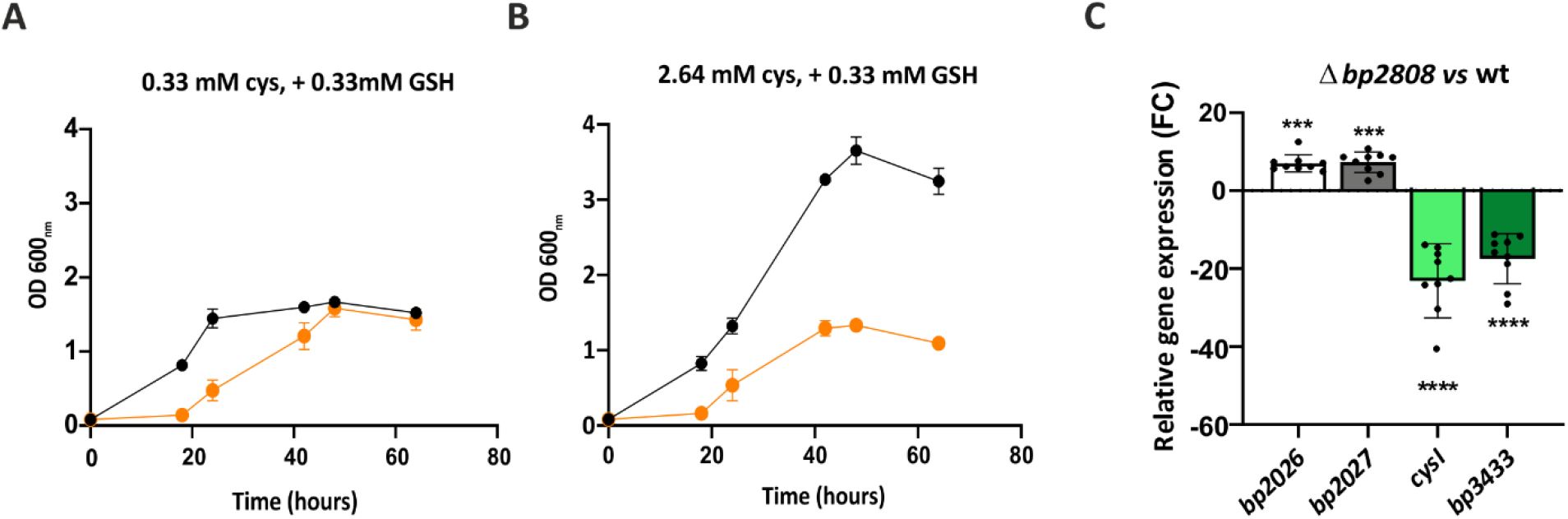
Growth and gene expression profiles of the Δ*bp2808* mutant in excess cysteine. The wt strain (black circles) and the Δ*bp2808* mutant (orange circles) were grown in standard SS medium (**A**) and in standard SS medium containing 2.64 mM cysteine (**B**). Data represent the mean optical density (OD_600_) ± SD from three independent biological replicates. (**C**) The expression of *bp2026, bp2027, cysI*, and *bp3433* genes was determined in biological triplicates of the wt strain and Δ*bp2808* mutant harvested 30 min after addition of 2.64 mM cysteine. Gene expression was compared between the Δ*bp2808* and wt cells. RT-qPCR analysis of each sample was performed in technical triplicate. The graphs show all individual data points, mean values, and error bars representing SDs. Statistical significance was calculated using one-way ANOVA (Dunnett’s multiple comparison test): (***) *P*-value < 0.0005 and (****) *P*-value < 0.0001.

Impaired growth of the Δ*bp2808* mutant in the presence of cysteine and sulfite suggested that the BP2808 exporter is required for sulfur homeostasis. Furthermore, its expression was highly induced by excess cysteine compared to other sulfur-related exporters such as BP2026 and BP2027 (Fig. 3). Thus, we examined whether and how *B. pertussis* compensates for the absence of the BP2808 exporter. We hypothesized that in the presence of excess sulfur, *B. pertussis* cells overexpress a), other exporter genes, such as *bp2026* and *bp2027*, or b), genes encoding enzymes that are required for recycling of cysteine metabolites such as sulfite reductase CysI (BP3432), or for the conversion of sulfite to sulfate such as sulfite oxidoreductase (BP3433). Therefore, we compared the expression of the *bp2026, bp2027, cysI*, and *bp3433* genes between the wt and Δ*bp2808* strains in the presence of excess cysteine. The wt and Δ*bp2808* cells were grown overnight in SS medium (0.32 mM GSH only) and then treated for 30 min with 2.64 mM cysteine (Fig. 6C). Total RNA isolated from harvested cells was used for quantitative PCR. Compared to the wt strain, the expression of both exporter genes was significantly induced in the Δ*bp2808* mutant, suggesting that these exporter genes compensate for the absence of the *bp2808* gene. In contrast, the expression of the *cysI* and *bp3433* genes was strongly reduced.

### The lack of *cdo* genes results in reduced secretion of pertussis toxin and attenuated virulence

Previously, it was shown that cysteine and glutathione affect the secretion of pertussis toxin (PT) by *B. pertussis* (19). PT is one of the major virulence factors (20) and consists of five distinct subunits (S1 to S5). After PT assembles in the periplasm, the holotoxin is secreted by the type IV secretion system (21). Recently, we showed that the strain lacking two *cdo* genes displays reduced cytotoxicity toward human macrophages (16). Therefore, we asked whether the reduced cytotoxicity could result from impaired secretion of PT caused by altered cysteine metabolism. The wt and Δ*bp2871*Δ*bp3011* strains were grown in standard SS medium and in SS medium supplemented with either 1.32 mM cysteine or 1.32 mM GSH, and the amounts of the S1 subunit of pertussis toxin (PT_S1_) were determined in pelleted cells and filtered culture supernatants using immunoblotting. Western blot analysis revealed that regardless of the source of cysteine, pelleted wt and mutant cells contained similar amounts of PT_S1_ protein (Fig. 7A). However, secretion of PT_S1_ protein by the mutant strain was strongly reduced in standard SS medium and SS medium supplemented with 1.32 mM cysteine compared to the wt strain. Interestingly, in the presence of 1.32 mM GSH, the mutant secreted similar amounts of PT_S1_ protein as the wt strain. Apparently, the lack of *cdo* genes resulted in impaired secretion of the PT in the presence of cysteine, while this defect was absent in the presence of GSH. PT contains 11 intramolecular disulfide bonds, and its proper folding and secretion also depend on the Dsb family of enzymes, DsbA and DsbC, respectively (19, 22). We compared the expression of *dsbA* and *dsbC* genes between the Δ*bp2871*Δ*bp3011* mutant and wt cells grown in the presence of 1.32 mM cysteine (no GSH). The expression of both *dsb* genes was significantly increased in the *cdo* mutant compared to the wt strain, while the expression of the *ptxA* gene encoding the PT_S1_ subunit was not affected (Fig. 7B). Since the secretion of PT was efficient in the presence of 1.32 mM GSH, we tested the expression of *ptxA, dsbA* and *dsbC* genes also under these conditions. In contrast to excess cysteine, the excess GSH did not significantly affect the expression of *dsb* genes (Fig. 7B). Collectively, these results suggest that the *cdo* mutant exposed to excess cysteine experiences disulfide stress, which results in impaired secretion of the PT.

**FIG 7.**
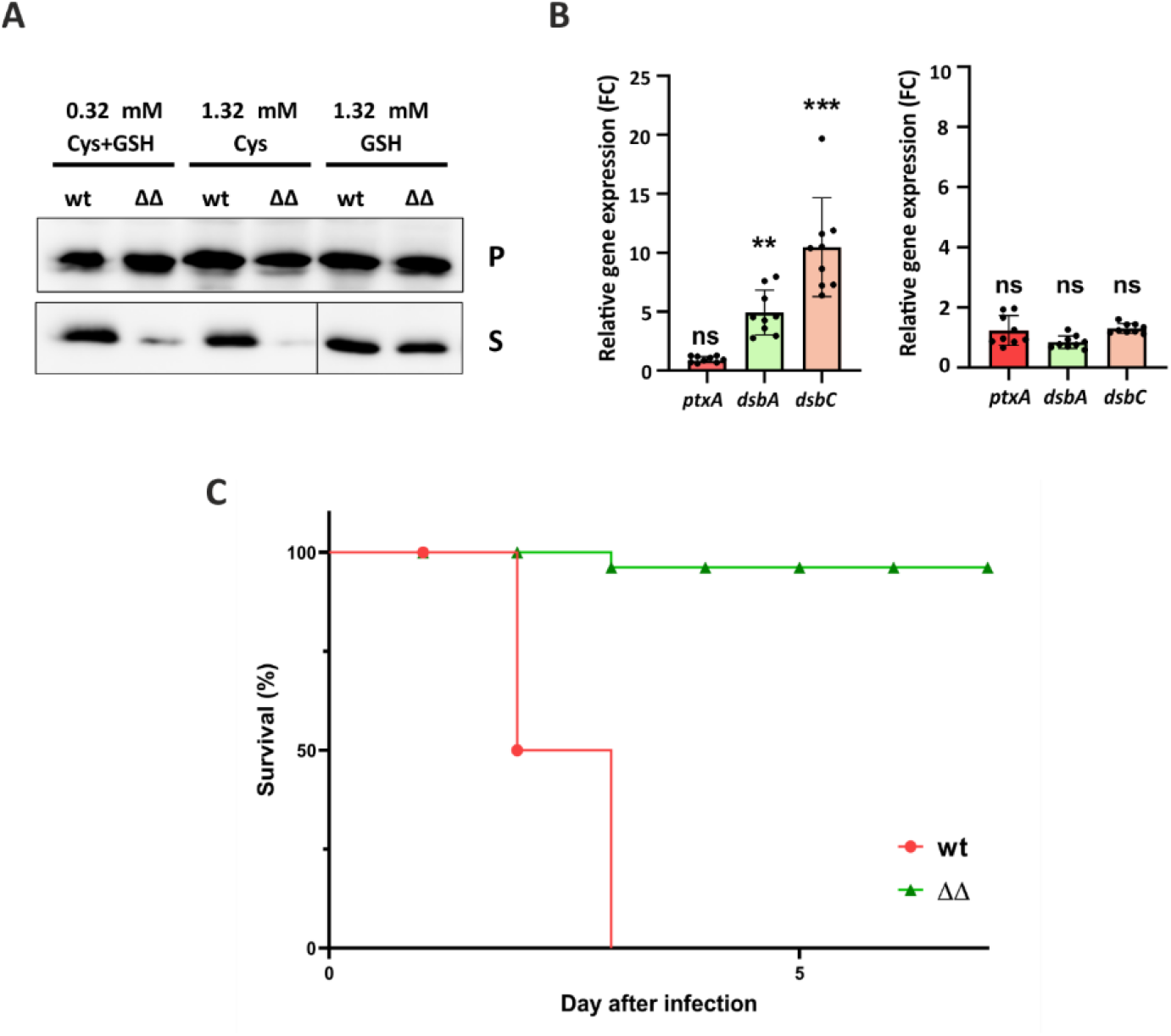
The absence of two *cdo* genes affects secretion of PT_S1_ and the virulence of *B. pertussis*. (**A**) The wt strain (wt) and the Δ*bp2871*Δ*bp3011* mutant (ΔΔ) were grown in standard SS medium (0.32 mM cysteine, 0.32 mM GSH) and in SS medium supplemented with either 1.32 mM cysteine or 1.32 mM GSH. Samples of pelleted cells (P) equivalent to 0.1 OD_600_ unit and precipitated supernatant proteins (S) equivalent to 1 OD_600_ unit were separated on a 12.5% SDS-PAGE and analyzed by immunoblotting using antibodies against the S1 subunit of pertussis toxin. Only the relevant parts of the membranes are shown. (**B**) The expression of *ptxA, dsbA*, and *dsbC* genes was determined in biological triplicates of the wt strain and Δ*bp2871*Δ*bp3011* mutant cells grown in the presence of 1.32 mM cysteine (no GSH; left plot) and 1.32 mM GSH (no cysteine; right plot). Gene expression was compared between Δ*bp2871*Δ*bp3011* and wt cells. RT-qPCR analysis of each sample was performed in technical triplicate. The graphs show all individual data points, mean values, and error bars representing SDs. Statistical significance was calculated using one-way ANOVA (Dunnett’s multiple comparison test): ns, not significant; (**) *P*-value < 0.005; (***) *P*-value < 0.0005. (**C**) Kaplan-Meier survival curves of mice infected with *B. pertussis*. Six mice per group were intranasally infected with 1.5 × 10^8^ of wt (wt) or Δ*bp2871*Δ*bp3011* mutant (ΔΔ), and the survival of mice was monitored for one week. The plot was generated using GraphPad Prism.

To analyze the impact of two *cdo* genes on *B. pertussis* virulence *in vivo*, we used a mouse intranasal model of infection. Two groups of six BALB/c mice were intranasally challenged with 1.5 × 10^8^ cells per mouse of either the wt or Δ*bp2871*Δ*bp3011* strains, and their survival was monitored for seven days. While the wt bacteria caused the death of all infected mice within three days, the same dose of Δ*bp2871*Δ*bp3011* bacteria killed only one mouse (Fig. 7C). Thus, deletion of the two *cdo* genes resulted in a substantial reduction of *B. pertussis* virulence in the mouse infection model.

## Discussion

To date, sulfur metabolism and its role in *B. pertussis* fitness and virulence have not been studied systematically. Pathogens often evolve streamlined metabolism, marked by the loss of redundant biosynthetic pathways and increased reliance on host-derived nutrients (23). A previous genomic study revealed that *B. pertussis* decayed its sulfate assimilation pathway, as several genes involved in sulfate transport and assimilation were pseudogenized or lost (24). Consequently, sulfate cannot serve as the primary inorganic sulfur source for cysteine biosynthesis, and *B. pertussis* is auxotrophic for cysteine (25). Amino acid auxotrophies are relatively common in bacterial pathogens and their auxotrophy must be compensated for by the nutritional background of the host (26-28). Cysteine auxotrophy is relatively frequent, especially among intracellular pathogens, and may reflect the high consumption of energy and reducing power linked to sulfate assimilation (27-29). For example, pathogens such as *Listeria monocytogenes, Legionella pneumophila*, and *Francisella tularensis* are cysteine auxotrophs due to incomplete sulfate assimilation pathways (30-32). Similar to these pathogens, *B. pertussis* cells must acquire cysteine from the host. We suggest that the decay of the sulfate assimilation pathway and cysteine auxotrophy can be considered part of a genome streamlining strategy that assists *B. pertussis* in pathoadaptation to its human host. Additionally, the sulfate metabolism of *B. pertussis* appears to be further simplified, for example, genes encoding the *O*-acetylserine (thiol)-lyase A and cystathionine γ-synthase, required for synthesis of cysteine from sulfide and methionine, respectively, are missing. Consequently, methionine does not support the growth of *B. pertussis* as a sole source of sulfur (33).

*B. pertussis* is an extracellular pathogen that colonizes the ciliated epithelium of the host upper respiratory tract. However, non-opsonized *B. pertussis* cells can, at least in part, survive phagocytic killing and proliferate in macrophages (34, 35). Recently, our transcriptomic profiling of intracellular *B. pertussis* cells revealed extensive rewiring of sulfur metabolism during infection of human macrophages (16). We observed that, as a part of adaptation to the intramacrophage environment, genes involved in cysteine catabolism and export were strongly induced. Importantly, our *in vitro* data largely recapitulate the results obtained with intracellular *B. pertussis* cells and thus help to elucidate the nature of the pathogen’s adaptive processes within the phagosome. For example, we show that the expression of both *cdo* genes *in vitro* can be induced by increased amounts of cysteine, suggesting that upon internalization, *B. pertussis* cells are exposed to excess cysteine. Given the reduced expression of the cysteine importer gene during infection (16) and the high content of glutathione (1-10 mM) in the cytoplasm of human cells (36) and in the alveolar epithelial lining fluid (37), we speculate that, similar to *F. tularensis* (30), *B. pertussis* predominantly uses glutathione as an exogenous source of cysteine. In support, *B. pertussis* encodes at least six ABC transporters with specificity for di- and oligopeptides that are homologous to glutathione ABC importer GsiABCD of *E. coli* (*e*.*g. bp2394*-*bp2397*), out of which five are significantly upregulated in intracellular bacteria (16).

The expression of the *bp2808* gene, which encodes the putative YeiH sulfate exporter, is highly induced upon addition of cysteine and is also significantly induced in internalized bacteria. Cysteine is metabolized by CDOs to cysteinesulfinic acid, which can be further metabolized to pyruvate and sulfite. Sulfite is a toxic and potent reducing agent and must be either exported or converted to sulfate or sulfide. Our data did not determine whether BP2808 exports sulfite and/or sulfate; however, growth of the mutant lacking the *bp2808* gene was strongly impaired by sulfite. Therefore, we hypothesize that the BP2808 protein is primarily required for sulfite export during cysteine-induced stress. Additionally, we show that in the absence of the BP2808 exporter, excess cysteine induces expression of the putative cysteine (*bp2026*) and sulfite (*bp2027*) exporter genes. We presume that increased expression of *bp2026* and *bp2027* compensates for the absence of the *bp2808* gene and enables the pathogen to cope with excess cysteine. In contrast, genes involved in sulfite metabolism, such as sulfite reductase (*cysI*) and sulfite oxidoreductase (*bp3433*), are strongly downregulated upon cysteine shock. Sulfite reductase CysI converts sulfite to sulfide, which can then be used with *O*-acetyl-L-serine to produce cysteine. This reaction is catalyzed by *O*-acetylserine (thiol)-lyase A, encoded by the *cysK* gene (4), which is, however, missing in *B. pertussis*. Thus, producing sulfide by CysI seems biologically meaningless and could lead to accumulation of sulfide, which can be toxic in higher amounts (38). Likewise, since the sulfate assimilation pathway is not functional in *B. pertussis*, conversion of sulfite to sulfate by sulfite oxidoreductase BP3433 may not provide much benefit. These circumstances most likely explain the strongly reduced expression of the *cysI* and *bp3433* genes and suggest that *B. pertussis* relies on exporting cysteine metabolites rather than recycling them, thereby limiting the levels of inorganic sulfur within the cell.

One of the most important findings of our study is the critical role of CDO enzymes in the fitness and virulence of a strictly human bacterial pathogen. These mononuclear non-heme iron enzymes were previously thought to be present only in the metazoan and fungal kingdoms (39). In mammals, CDOs contribute to cysteine metabolism and balance, and mutants lacking CDO display severe defects (40, 41). Later, CDOs were also discovered in prokaryotes, as Dominy and co-workers showed that cysteine can be an authentic substrate for CDOs in several eubacteria (42). Nevertheless, the importance of CDOs for sulfur metabolism and the fitness of bacterial pathogens was not fully recognized. Although the thiol group in cysteine can be a potential source of toxicity (6, 43), our data show that *B. pertussis* can tolerate and metabolize millimolar concentrations of cysteine, and that this ability largely depends on the activity of CDO enzymes and GSH. This suggests that the decay of the sulfate assimilation pathway in *B. pertussis* is not due to potential cysteine toxicity but rather represents an adaptation to its human host. While eukaryotic CDO enzymes contain close to the active site a thioether crosslink between the sulfur of cysteine (C93) and the carbon of tyrosine (Y157), in most bacterial CDOs the position equivalent to cysteine is replaced by glycine (44-46). Nevertheless, some CDOs in several Gram-negative pathogenic bacteria have cysteine analogous to C93 (39). CDOs with a crosslink exhibit higher activity and stability in the presence of millimolar concentrations of cysteine (44). Furthermore, the optimal pH of CDOs possessing C93 is approximately 6.6, while the C93G variant has a pH optimum of 7.3 (45). Given the millimolar concentrations of GSH in mammalian cells and the acidic pH within the phagosome, it could be expected that the bacterial CDO enzyme containing cysteine would be more active within the host than its glycine variant. Strikingly, both *B. pertussis* CDO enzymes, BP2871 and BP3011, contain cysteine near the active site and are highly induced in phagocytosed cells. Thus, we speculate that *B. pertussis* is capable of cysteine-tyrosine crosslink formation in both CDO enzymes, which improves its survival within a mammalian host. Supporting this, our previous (16) and current data show that CDOs are required for the fitness and virulence of *B. pertussis in vitro, in cellulo*, and in the mouse model *in vivo*.

Our data also provide a possible explanation for the reduced virulence of the Δ*bp2871*Δ*bp3011* mutant in the mouse model of infection. Although we realize that the lack of *cdo* genes may have several effects, our data show that the absence of *cdo* genes affects the secretion of pertussis toxin, one of the major virulence factors of *B. pertussis* (20). Even at standard cysteine concentrations, secretion of pertussis toxin by the *cdo* mutant was reduced compared to the wt strain and this defect was further exacerbated by increased cysteine levels. Pertussis toxin (subunits S1 to S5) contains 11 intramolecular disulfide bonds formed by periplasmic Dsb proteins of the thiol oxidoreductase family and proper formation of these disulfide bonds is required for folding and secretion of pertussis toxin (22). Later, it was shown that secretion of pertussis toxin is also sensitive to the redox state of the periplasmic space, as the addition of glutathione, but not cysteine, improved toxin secretion (19). In contrast to this report, our study revealed that the wt strain, but not the *cdo* mutant, secreted similar amounts of toxin in media supplemented with either glutathione or cysteine. This result suggests that the redox buffering capacity of cysteine is sufficient for toxin secretion unless cysteine levels are imbalanced due to the absence of CDOs. We hypothesize that excess cysteine, combined with a lack of CDO activity, leads to cysteine accumulation in the periplasmic space, resulting in disruption of redox homeostasis and aberrant formation of disulfide bonds, as indicated by increased expression of the *dsbA* and *dsbC* genes.

In conclusion, our combined *in cellulo* and *in vitro* data suggest that sulfur metabolism of *B. pertussis* within the host is relatively simple (Fig. 8). Cysteine, most likely derived from GSH provided by the host, is used for cellular functions. Because of its potential toxicity, excess cysteine is scavenged by the two CDOs, BP2871 and BP3011, and eventually converted to pyruvate, which serves as a source of energy and toxic sulfite, which is exported by the sulfite exporter BP2808. In this scenario, *B. pertussis* cells lacking either CDO enzymes or the BP2808 exporter experience cysteine-induced stress that disrupts redox balance and, consequently, physiological fitness and virulence. These observations suggest that targeting the import of host-derived GSH and the export of sulfite could serve as an antimicrobial therapy.

**FIG 8.**
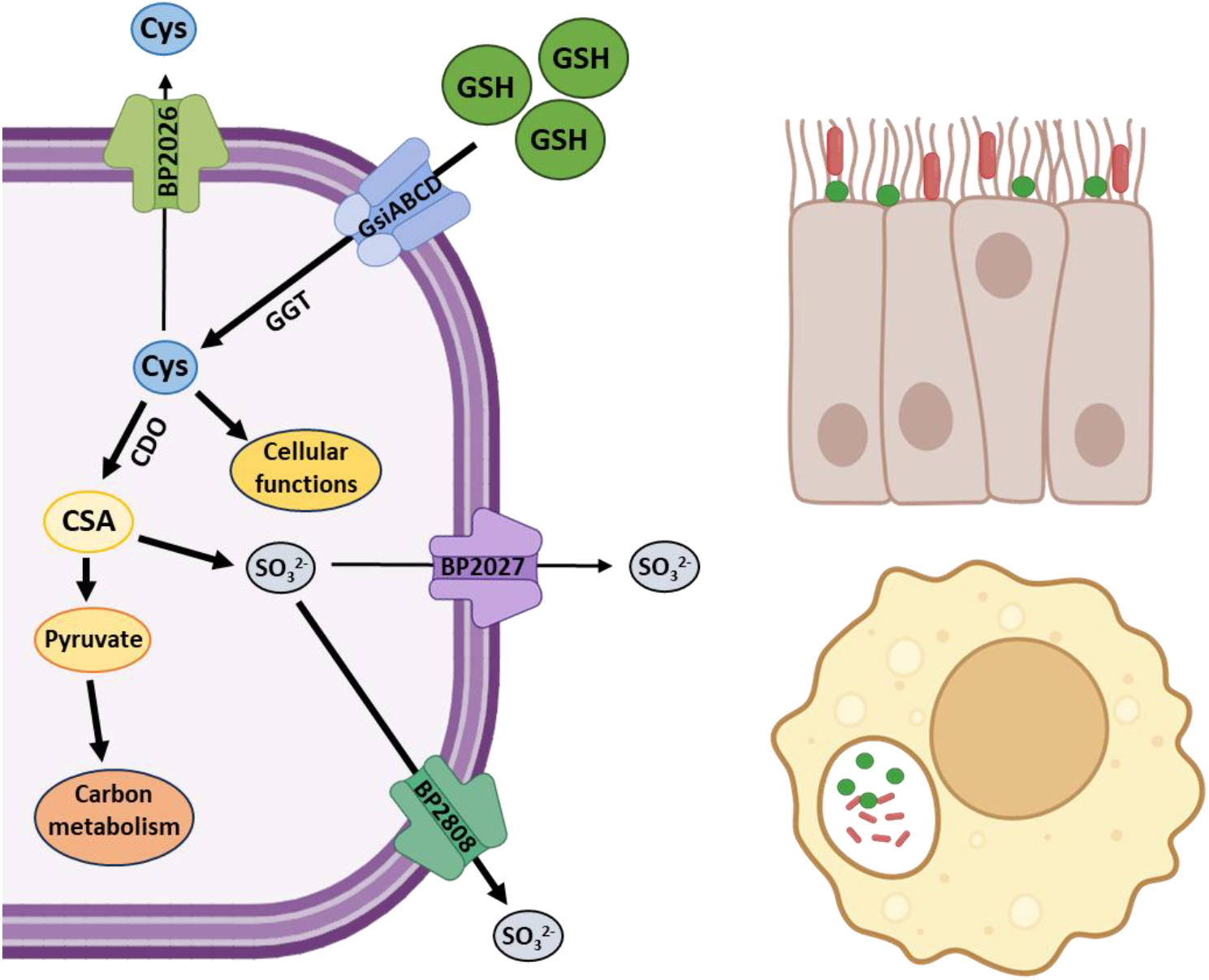
Sulfur metabolism is streamlined in *B. pertussis* following infection of the host. The schematic diagram shows sulfur metabolism in *B. pertussis* cells during infection. Eukaryotic cells, such as ciliated epithelial cells or macrophages contain millimolar concentrations of glutathione (GSH, green marbles). GSH serves as a cysteine source for *B. pertussis* cells (purple rods) that colonize ciliated respiratory epithelium or are internalized by phagocytes. *B. pertussis* imports GSH via several ABC transporters of the Gsi family and converts it to cysteine through periplasmic or cytosolic γ-glutamyl transpeptidase (GGT). Cysteine is then used for cellular functions such as protein synthesis, Fe-S clusters production, and coenzymes biosynthesis. Part of the cysteine is converted to cysteine sulfinic acid (CSA) by the cysteine dioxygenases (CDO) BP2871 and BP3011. CSA is then catabolized to pyruvate, which is used as an energy source, and sulfite (SO_3_ ^2-^), which is transported by the sulfite exporter BP2808. Excess cysteine and sulfite can alternatively be exported by transporters BP2026 and BP2027, respectively. Thick arrows indicate major pathways of cysteine metabolism based on our *in vitro* and *in cellulo* data; thin arrows indicate alternate pathways.

## Materials and methods

### Bacterial cultures and growth conditions

*Bordetella pertussis* strain Tohama I and its derivatives were grown on charcoal agar (Oxoid, CM0119, United Kingdom) plates for 3 days at 37 °C in a humidified incubator (5 % CO_2_). For planktonic cultures, bacteria were grown in standard Stainer-Scholte (SS) medium (25), *i*.*e*. without casamino acids and cyclodextrin, at 37 °C. Standard SS medium contains 0.33 mM cysteine and 0.32 mM GSH. To study the effects of cysteine and GSH separately, in some experiments the SS medium contained only cysteine (1.32 mM or 2.64 mM) or only GSH (0.32 mM or 1.32 mM), as specified in the figure legends.

### Construction of the mutants

The deletions of the *bp0598* and *bp2808* genes were introduced into the chromosome of the *B. pertussis* Tohama I strain, referred to as the wild-type (wt) strain in this study, as described previously (47, 48). Briefly, to generate a deletion mutant, two DNA fragments of approximately 700 bp flanking the corresponding gene were amplified from the upstream region (ending with the initiation codon followed by an *Nhe*I restriction site) and the downstream region (beginning with the *NheI* site followed by the stop codon). The resulting PCR products were ligated over the *Nhe*I site, and the ligation mixture was used to amplify an approximately 1.4 kb PCR product in which the start and stop codons, separated by the *Nhe*I site, created a markerless in-frame deletion. For both constructs, the final PCR products were ligated into the allelic exchange plasmid pSS4245 (49) and cloned in *E. coli* XL-Blue cells. The resulting recombinant plasmids were transformed into the *E. coli* SM10 λ-pir strain and transferred into the *B. pertussis* Tohama I strain by conjugation. After two recombination events, the strain carrying the mutation was obtained and verified by sequencing of the amplified PCR product covering the adjacent chromosomal regions. The primers used for generation of the mutants are listed in Table S1.

### RNA isolation and RT-qPCR

Gene expression profiles were determined as recently described (48). Total RNA was isolated from lysed cells using TRI reagent (Sigma) according to manufacturer’s protocol. Removal of DNA was achieved by treatment of samples with DNase I Solution (1 unit/μL, Sigma-Aldrich) for 30 minutes at 37 °C. RNA concentration was measured using Nanodrop 2000 instrument (Thermo Fisher Scientific). Total RNA (1 µg) was reverse transcribed into cDNA using random hexamers and M-MLV reverse transcriptase (Promega) at 37 °C. Reactions (25-μl) were performed in at least three technical replicates per sample on a Bio-Rad CFX96 instrument using HOT FIREPol® EvaGreen® qPCR Mix Plus (Solis Biodyne), 4 pmoles of each primer and 2 µl of 30-fold diluted cDNA in a 20-μl reaction was pipetted into 96-well plates. Each reaction consisted of an initial step at 95 °C for 12 min, followed by 40 cycles of 95 °C for 15 s, 62 °C for 20 s, and 72 °C for 20 s, followed by melting curve recording. The *rpoB* gene was used as the reference gene and relative gene expression was determined using the delta-delta C_t_ method (50). The sequences of primers used for RT-qPCR are listed in Table S1.

### Western blot analysis

The amounts of S1 subunit of pertussis toxin in lysates and supernatants of *B. pertussis* cultures were determined as previously described (48). Samples of pelleted cells equivalent to 0.1 OD_600_ unit or secreted proteins precipitated from filtered culture supernatants equivalent to 1.0 OD_600_ unit were separated on 12.5% SDS-polyacrylamide gels and transferred onto a nitrocellulose membrane using the Trans-Blot® Turbo™ system (Bio-Rad). Membranes were blocked with 5% skim milk and probed with polyclonal antibodies raised against the S1 subunit of pertussis toxin (1:20 000), followed by incubation with anti-mouse IgG antibodies conjugated with horseradish peroxidase (Cell Signaling Technology, Inc.) at a 1:10 000 dilution. Antibody-antigen complexes were visualized using SuperSignal West Femto chemiluminescent substrate (Thermo) according to the standard protocol on the G:Box Chemi XRQ device (Syngene).

### Mouse infection

Six-week-old Balb/c mice were obtained from Charles River Laboratories and maintained under specific pathogen-free conditions. *B. pertussis* wt and Δ*bp2871*Δ*bp3011* strains were grown in SS medium to the exponential phase of growth at 37 °C, OD_600_ values were measured, and bacterial cells were appropriately diluted in PBS to same density. Groups of six mice were anesthetized by intraperitoneal injection of ketamine (80 mg/kg) and xylazine (8 mg/kg) and the wt and Δ*bp2871*Δ*bp3011* cells were administered intranasally in suspensions of 50 μL PBS containing 1.5 × 10^8^ bacteria/mouse. Survival of mice was monitored twice a day for one week. Results are expressed as the percentage of surviving mice and are representative of two independent experiments. Log-rank tests were used to determine the significance of the differences in survival between the groups of mice, *p* values (at 95 % confidence intervals) and Kaplan-Meier plots were generated using GraphPad Prism (version 8.4.3).

## Ethics statement

Animal experiments were approved by the Animal Welfare Committee of the Institute of Molecular Genetics of the Czech Academy of Sciences in Prague, Czech Republic. Handling of animals was performed according to the *Guidelines for the Care and Use of Laboratory Animals*, the Act of the Czech National Assembly, Collection of Laws no. 246/1992. Permission no. 1081/2025 was issued by the Animal Welfare Committee of the Institute of Molecular Genetics of the Czech Academy of Sciences in Prague.

## Acknowledgments

We thank Ondřej Staněk for providing polyclonal antibodies raised against the S1 subunit of pertussis toxin. Computational resources were provided by the e-INFRA CZ project (ID:90254), supported by the Ministry of Education, Youth and Sports of the Czech Republic.

## Author contributions

Conceptualization: Argha Saha, Branislav Večerek; Methodology: Argha Saha, Denisa Petráčková, Jana Holubová; Investigation: Argha Saha, Denisa Petráčková, Jana Holubová; Data analysis: Argha Saha, Denisa Petráčková, Branislav Večerek; Data curation and visualization: Argha Saha, Denisa Petráčková; Funding acquisition: Branislav Večerek; Writing-original draft: Argha Saha, Branislav Večerek; Writing-review and editing: Argha Saha, Branislav Večerek

## Data and materials availability

All data needed to evaluate and reproduce the results in the article are present in the article and/or the Supplementary Materials. All the strains generated for this study are available from the corresponding author upon request.

## Supporting information caption

**Fig. S1 *Bordetella pertussis* has decayed or lost several genes in the sulfate assimilation pathway.** The schematic shows the sulfate assimilation pathway in bacteria. Genes that were decayed, such as *cysUW, cysN*, and *cysH*, or lost, such as *cysC* and *cysK*, in *B. pertussis* are shown in orange and red, respectively.

**Fig. S2 The homology of the putative GshA protein BP0598 to other GshA proteins.** (A) Sequence alignment of the putative glutamate-cysteine ligase (GshA) BP0598 (Uniprot ID Q7VS51) with other GshA proteins from *Ralstonia nicotinae* (Q8XU95), *Cupravidus metallidurans* (Q1LHL9), *Escherichia coli* (p77213), *Salmonella Typhimurium* (O68838), *Cronobacter sakazakii* (A7MJ27), *Pseudomonas aeruginosa*(Q8ZNK7), *Achromobacter denitrificans* (A0A3R9G2Y1).

**Fig. S3 The exporter BP2808 is homologous to the YeiH family of exporters.** (A) Sequence alignment of the YeiH exporter BP2808 (Uniprot ID Q7VV78) with other YeiH family exporters from *Achromobacter denitricans* (A0ABZ3G5K1), *Burkholderia stabilis* (A0AAJ5T2Q2), *Cronobacter sakazakii* (A7MLJ6), *Escherichia coli* (P62723), *Pseudomonas aeruginosa* (Q9HTI1), *Salmonella Typhimurium* (Q8ZNK7), and *Serratia rubidaea* (A0A448SJY0). Amino acid residues (Leu48, Ala148, Glu208, Val209, Ala210) predicted to form the ligand binding site are shown with red boxes (B) Localization of amino acid residues (in red) predicted to participate in the ligand binding site on the surface of BP2808 exporter. Visualization and prediction were performed using PrankWeb, a computational tool for predicting ligand binding sites (https://elixir-europe.org/services/prankweb).

**Table S1. Primers used in this study**

